# Liquid-liquid extraction of lipidated peptides for direct identification of lipidation sites

**DOI:** 10.1101/2023.05.25.542030

**Authors:** Kazuya Tsumagari, Yosuke Isobe, Yasushi Ishihama, Jun Seita, Makoto Arita, Koshi Imami

**Affiliations:** Proteome Homeostasis Research Unit, RIKEN Center for Integrative Medical Sciences, Tsurumi-ku, Yokohama, Kanagawa 230-0045, Japan; Laboratorhy for Metabolomics, RIKEN Center for Integrative Medical Sciences, Tsurumi-ku, Yokohama, Kanagawa 230-0045, Japan; Laboratorhy for Integrative Genomics, RIKEN Center for Integrative Medical Sciences, Tsurumi-ku, Yokohama, Kanagawa 230-0045, Japan; Division of Physiological Chemistry and Metabolism, Graduate School of Pharmaceutical Sciences, Keio University, Minato-ku, Tokyo 105-8512, Japan; Department of Molecular Systems Bioanalysis, Graduate School of Pharmaceutical Sciences, Kyoto University, Kyoto 606-8501, Japan; Laboratory of Clinical and Analytical Chemistry, National Institute of Biomedical Innovation, Health and Nutrition, Ibaraki, Osaka 567-0085, Japan; Cellular and Molecular Epigenetics Laboratory, Graduate School of Medical Life Science, Yokohama City University, Tsurumi-ku, Yokohama, Kanagawa 230-0045, Japan; WPI-Bio2Q, Keio University, Japan

## Abstract

Proteins can be modified by lipids in various ways, for example by myristoylation, palmitoylation, farnesylation, and geranylgeranylation—these processes are collectively referred to as lipidation. Current chemical proteomics using alkyne lipids has enabled the identification of lipidated protein candidates but does not identify endogenous lipidation sites and is not readily applicable to *in vivo* systems. Here, we introduce a proteomic methodology for global analyses of endogenous lipidation sites that combines liquid-liquid extraction of hydrophobic lipidated peptides with liquid chromatography-tandem mass spectrometry using a gradient program of acetonitrile in the high concentration range. We applied this method to explore lipidation sites in HeLa cells, and identified a total of 90 lipidation sites, including 75 protein N-terminal myristoylation sites, which is more than the number of high-confidence lipidated proteins identified by myristic acid analog-based chemical proteomics. Isolation of lipidated peptides from digests prepared with different proteases enabled the identification of different lipidated sites, extending the coverage. Moreover, our peptide-centric approach successfully identified dually modified peptides having myristoylation and palmitoylation. Finally, we analyzed *in vivo* myristoylation sites in mouse tissues and found that the lipidation profile is tissue-specific. This simple method (not requiring chemical labeling or affinity purification) should be a promising tool for global profiling of various protein lipidations.

## Introduction

Numerous proteins are covalently modified with a variety of lipid molecules, as exemplified by myristoylation of protein N-terminal glycine, and palmitoylation, farnesylation, and geranylgeranylation of cysteine^1^. These processes are collectively referred to as lipidation. In addition to these major lipidations, recent studies have revealed that other forms of lipids, such as cholesterol^1,2^ phosphoethanolamine^3,4^, and octanoic acid,^5^ can covalently modify proteins. Lipidation alters protein hydrophobicity, thereby affecting stability, localization, and interaction with other proteins^6,7^. For instance, N-terminal myristoylation of proto-oncogene tyrosine-protein kinase SRC (SRC) influences the protein folding and thereby promotes a switch to the active conformation^8^. Elevated myristoylation levels of SRC can cause cancer^8^, suggesting that lipidation may be involved in disease mechanisms. While protein lipidation is one of the key modifications that drive many aspects of biological processes, it has been less studied than other modifications such as phosphorylation, possibly due to the technical difficulties in analyzing low-abundance hydrophobic peptides, as well as the structural diversity of lipid modifications.

Proteomics using nanoscale liquid chromatography–tandem mass spectrometry (nanoLC/MS/MS) is a powerful approach, particularly in studies of post- or co-translational modifications of proteins. For large-scale analysis of modification sites of interest, modified peptides are biochemically enriched from protein digests using an affinity matrix, such as immobilized antibodies, since their abundance is relatively low compared to unmodified peptides. Such peptide-level enrichment is the key to pinpointing modification sites with regulatory functions, but enrichment of lipidated peptides is challenging. Until now, lipidated proteins have been analyzed using a chemical proteomic approach, in which proteins are metabolically labeled with lipid-mimicking chemical probes and isolated by affinity purification via click chemistry, followed by identification of unmodified peptides from digested lipidated proteins^9^. For instance, Thinon et al. performed protein N-terminal myristoylome analysis of HeLa cells using clickable myristic acid analogs, and identified 70 co-translationally myristoylated proteins with high confidence^10^. Similarly, chemical proteomic identification of palmitoylated proteins^11^ and farnesylated and geranylgeranylated proteins^12^ has been reported. While these chemical proteomic approaches are effective to profile the lipidated proteome, they have several limitations. First, site-level information is often lost. Second, chemical probes may not capture endogenous lipid modifications due to structural differences from endogenous lipids. Third, the limited availability of chemical probes limits the ability to analyze diverse lipid modifications. Fourth, the metabolic labeling approach is not readily applicable *in vivo*, such as in animal tissues. Thus, a new strategy that overcomes these limitations is now needed, especially because advanced lipidomics technologies have recently established that lipids in living organisms are more complex and diverse than previously believed^13,14^, raising the possibility of discovering novel protein modifications by lipids.

Herein, we introduce a proteomic approach for directly identifying protein lipidation sites. Lipidated peptides are isolated from protein digests by liquid-liquid extraction (LLE) and analyzed by nanoLC/MS/MS. We provide a proof-of-concept of this methodology targeting HeLa cells, and then describe a further application for quantitative analysis of *in vivo* lipidation sites in mouse tissues.

## Experimental Section

### Cell culture

HeLa cells were cultured in Dulbecco’s modified Eagle’s medium (FUJIFILM Wako, Osaka, Japan) containing 10 % FBS (Thermo Fisher Scientific, Waltham, MA) and 100 U/mL penicillin and 100 μg/mL streptomycin (FUJIFILM Wako). Cells were harvested, washed with PBS and stored at -80 °C until use.

### Mouse

Animal experimental procedures were approved by the Animal Care and Use Committee of RIKEN. A male C57BL/6J mouse was purchased from CLEA Japan Inc. (Tokyo, Japan), bred under specific pathogen-free conditions, and sacrificed at 12 weeks of age. Organs were quickly dissected, snap-frozen with liquid nitrogen, and stored at -80 °C until use.

### Protein extraction and digestion

HeLa cells were suspended in 8 M urea buffer including 100 mM Tris-HCl (pH 8.5), 10 mM tris(2-carboxyethyl)phosphine (TCEP; FUJIFILM Wako), 40 mM 2-chloroacetamide (CAA; FUJIFILM Wako), and 28.3 U of benzonase (Merck, Darmstadt, German) and agitated at room temperature for 30 min. Then, protein was extracted and denatured by sonication for 20 min on ice. The protein concentration was determined by means of bicinchoninic acid (BCA) assay (Thermo Fisher Scientific). The protein solution was diluted 5-fold with 50 mM ammonium bicarbonate (ABC; FUJIFILM Wako), and proteins were digested overnight with LysC (FUJIFILM Wako) and trypsin (sequence grade; Promega, Madison, WI) at the protein:enzyme ratio of 100:1 for each enzyme at 25 °C. When GluC (FUJIFILM Wako) or chymotrypsin (ROCHE, Basel, Switzerland) was used, the protein solution was diluted 10-fold with 50 mM phosphate buffer (pH 8.5; for GluC) or ABC (for chymotrypsin) and digested at the protein:enzyme ratio of 50:1. Digestion was halted by acidifying the mixture with trifluoroacetic acid (TFA; FUJIFILM Wako). The obtained digests were purified on MonoSpin C18 reversed-phase columns (GL Sciences, Tokyo, Japan).

Mouse organs were suspended in 5 % sodium dodecyl sulfate (SDS) buffer containing 100 mM Tris-HCl (pH 8.5), 10 mM TCEP, and 40 mM CAA and crushed using zirconia beads (Tomy, Tokyo, Japan) and TissueLyser (Qiagen, Hilden, German). Proteins were inactivated at 95 °C for 5 min, extracted by sonication for 20 min on ice, and purified by methanol-chloroform precipitation. The protein concentration was determined by means of BCA assay. Proteins were digested with trypsin and LysC and the peptides were purified as described above.

### Liquid-liquid extraction

All organic solvents used for LLE were obtained from FUJIFILM Wako. In most experiments, 100 μg of desalted protein digest was subjected to LLE. In the experiment comparing solvents (related to Figure 2), 50 μg of digest was subjected to LLE. Digests were dissolved in 50 μL of 0.5 % TFA and vigorously mixed with 65 μL of organic solvent for 10 min using a vortex mixer at room temperature. Following centrifugation at 18,000 *g* for 5 min at room temperature, 50 μL of the upper phase (organic phase) was obtained and evaporated using a SpeedVac (Thermo Fisher Scientific).

### NanoLC/MS/MS

A nanoLC/MS/MS system comprising an EASY-nLC 1200 (Thermo Fisher Scientific) and an Orbitrap Eclipse mass spectrometer (Thermo Fisher Scientific) was employed. The mobile phases consisted of (A) 0.1 % formic acid and (B) 0.1 % formic acid and 80 % acetonitrile (ACN). Peptides were loaded on a lab-made 15-18 cm fused-silica emitter (100 μm inner diameter) packed with ReproSil-Pur C18-AQ (1.9 μm, Dr. Maisch, Ammerbuch, Germany), and separated as follows: first, contaminating hydrophilic peptides were washed out with 38 % B for 10 min at the flow rate of 800 nL/min, and then remaining hydrophobic peptides were separated by applying a linear gradient for 43 min (38–70 % B over 30 min, 70–99 % B over 3 min, and 99 % B for 10 min) at the flow rate of 500 (solvent comparison) or 300 nL/min (HeLa cells and mouse organs). MS scanning was initiated 10 min after the start of LC. All spectra were obtained using the Orbitrap analyzer. MS1 scans were performed in the range of 375– 1500 *m/z* (resolution = 120,000, maximum injection time = “Auto”, and automatic gain control = “Standard”). For the subsequent MS/MS analysis, precursor ions were selected and isolated in a top-speed mode (cycle time = 3 sec and isolation window = 1.6 *m/z*) and activated by higher-energy collisional dissociation (normalized collision energy = 28). Column temperature was set to 50 °C. In the analyses of HeLa cells and mouse organs, FAIMSpro (Thermo Fisher Scientific) was employed, and the samples were analyzed in four runs with different compensation voltages (−30/-70; -40/-80; -50/-90; and -60/-100).

In the experiment comparing retention times of synthetic lipidated peptides with HeLa digest (related to Fig. S1), 200 ng of HeLa digest spiked with 2 pmol synthetic peptides was injected. In the experiment comparing organic solvents (related to Fig. 2), isolated peptides were dissolved in 10 μL of loading buffer consisting of 4 % ACN and 0.5 % TFA, and an aliquot of 5 μL was injected into the MS. In the HeLa cells and mouse organ profiling (related to Fig. 3, 4), isolated peptides from 100 μg digest were dissolved in 22 μL of the loading buffer, and a 5 μL aliquot was injected per measurement.

The proteomics data have been deposited to the ProteomeXchange Consortium via the jPOST partner repository^15^ with the dataset identifier PXD041499 (JPST002094).

### Data processing

In most experiments, LC/MS/MS raw data were processed using FragPipe (v.19.0) with the MSFragger search engine (v.3.7), Philosopher (v.4.6.0), and IonQuant (v.1.8.10)^16^. Database search was implemented against the UniprotKB/SwissProt (April 2022) human or mouse database with commonly observed contaminant proteins. Cysteine carbamidomethylation as a fixed modification, and methionine oxidation and acetylation on the protein N-terminus as variable modifications were set in all searches. For the identification of myristoylation and palmitoylation sites, myristoylation of protein N-terminal glycine (+210.19836) and palmitoylation of cysteine (+181.2082) were considered as variable modifications. For identification of farnesylation and geranylgeranylation sites, farnesylation (+161.18199) and geranylgeranylation (+229.24458) of protein C-terminal cysteine were set as variable modifications, and digest specificity was set to semi-specific, considering that these modifications occur followed by proteolytic cleavage and C-terminal capping by methylation. Minimal peptide length was set to 6 amino acids. Match between runs was enabled in the mouse organ profiling experiment. Other parameters remained at the default settings. False discovery rates were estimated by searching against a reversed decoy database and filtered for <1% at the peptide-spectrum match (PSM) and protein levels.

In the case of HeLa cell profiling (related to Fig. 3), MaxQuant (v.2.1.1.0)^17^ was utilized, since MaxQuant allows simultaneous analysis of peptides prepared by different digestive enzymes. The parameters described above were employed, and the other parameters remained at the default settings. False discovery rates were estimated by searching against a reversed decoy database and filtered for <1% at the PSM and protein levels.

For confident identification of lipidated peptides in addition to FDR control, only those that met at least one of the following criteria were accepted: 1) the lipidated site is reported in the UniProt database, 2) the site was identified in two or more unique peptides, 3) the site was indicated by a peptide identified from three or more MS/MS spectra in which three or more sequential b- or y-ions were observed. In the case of farnesylation and geranylgeranylation sites, in addition to the above criteria, only sites on the protein C-terminus with a C*AAX* (*A*, aliphatic amino acid; *X*, any amino acid) motif^6^ were accepted.

### Downstream analysis

A sequence logo was created using WebLogo (v.1.0)^18^. Data visualization, Pearson correlation calculation, and hierarchical clustering were performed using the R framework (v.4.1.3) with the basic functions and the ggplot2 (v.3.3.6) and pheatmap (v.1.0.12) packages.

## Results and Discussion

### Isolation of lipidated peptides by liquid-liquid extraction

For detection of lipidated peptides by nanoLC/MS/MS, we first evaluated the chromatographic behavior of six synthetic lipidated peptides (three N-terminally myristoylated peptides and three cysteine *S*-palmitoylated peptides) spiked into HeLa tryptic digests. As expected, the lipidated peptides were eluted in a higher ACN concentration range (>32% ACN) than many other peptides (Fig. S1), suggesting that a special gradient condition is needed for analysis of hydrophobic lipidated peptides. We, therefore, sought to analyze lipidated peptides more effectively by using the strategy shown in Fig. 1. Firstly, similar to the conventional shotgun proteomics workflow, proteins extracted from cells are reduced, alkylated, digested, and desalted. Secondly, highly hydrophobic lipidated peptides from the protein digest are separated using LLE, where lipidated peptides partition into low-polarity organic solvents. Thirdly, we employ nanoLC/MS/MS with an atypical gradient program that starts from 30 % ACN for detection of lipidated peptides (see the experimental section for the detailed gradient conditions).

**Figure 1.**
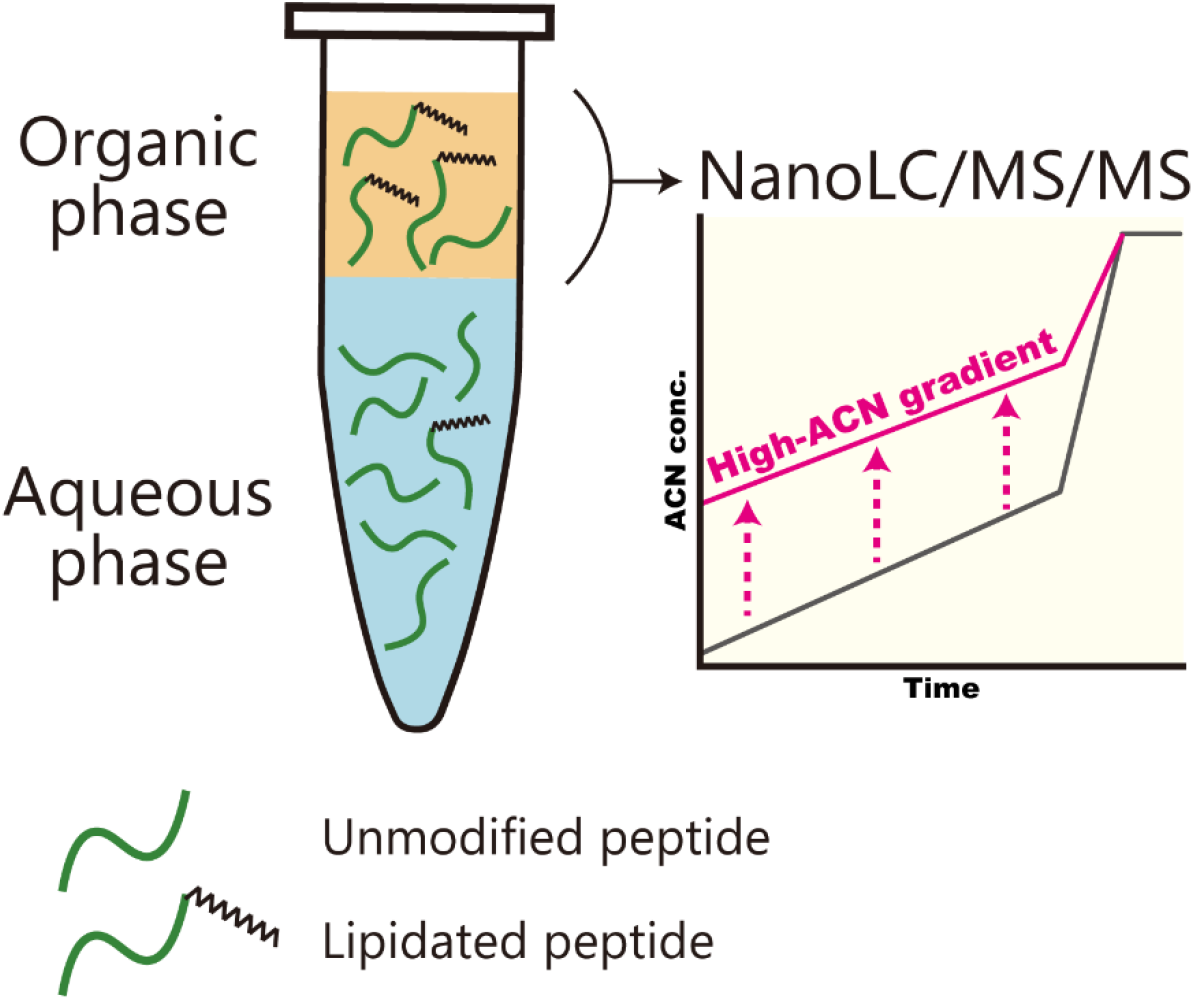
Scheme of extraction and analysis of lipidated peptides. Protein digests are prepared according to a conventional shotgun proteomics protocol. Hydrophobic lipidated peptides can partition into organic solvents and be isolated by liquid-liquid extraction. Isolated peptides are analyzed by nanoLC/MS/MS, focusing on the high acetonitrile (ACN) concentration range.

We first examined if peptides with N-terminal myristoylation at glycine or palmitoylation at cysteine residues are enriched using LLE. Previous studies by the Jensen’s group have shown that lipidated peptides can be isolated from protein digests by LLE^19,20^, but only ethyl acetate was tested as an organic solvent. We thus optimized LLE with various organic solvents, namely ethyl acetate, hexanol, heptanol, octanol, hexane, or toluene, using a trypsin/LysC digest of extracted HeLa cell proteins. More than 30 myristoylated peptides and several palmitoylated peptides were efficiently extracted by ethyl acetate, hexanol, heptanol, or octanol (Fig. 2A, S2). On the other hand, only a few myristoylated or palmitoylated peptides were extracted by hexane and toluene. We also examined if peptides with C-terminal farnesylation or geranylgeranylation at a cysteine residue can be enriched by an additional semi-specific search, since these modifications occur followed by proteolytic cleavage^6^. As a result, one or two farnesylated or geranylgeranylated peptides were identified using hexanol, heptanol, octanol, but no sites were identified using hexane or toluene (Fig. 2B, S2). Of note, we reanalyzed a publicly available deep HeLa proteome dataset in which more than 10,000 proteins were identified with the use of high pH reversed-phase fractionation mainly in the range of 5–35 % ACN^21^. However, no myristoylated peptides were identified despite the extensive fractionation of peptides (Fig. S3), highlighting the importance of isolating lipidated peptides then applying nanoLC/MS/MS with a high ACN gradient program. Moreover, we measured HeLa cell digest separated by a linear gradient ranging from 4 to 80 % ACN without LLE, but fewer myristoylated peptides were identified than were identified using LLE together with the high ACN gradient (Fig. S3). Thus, we concluded that both enriching lipidated peptides by LLE and analyzing peptides in the higher ACN concentration range are critical for the efficient detection of lipidated peptides. We decided to use hexanol and heptanol for further analyses as they provided the highest identification number of lipidated peptides.

**Figure 2.**
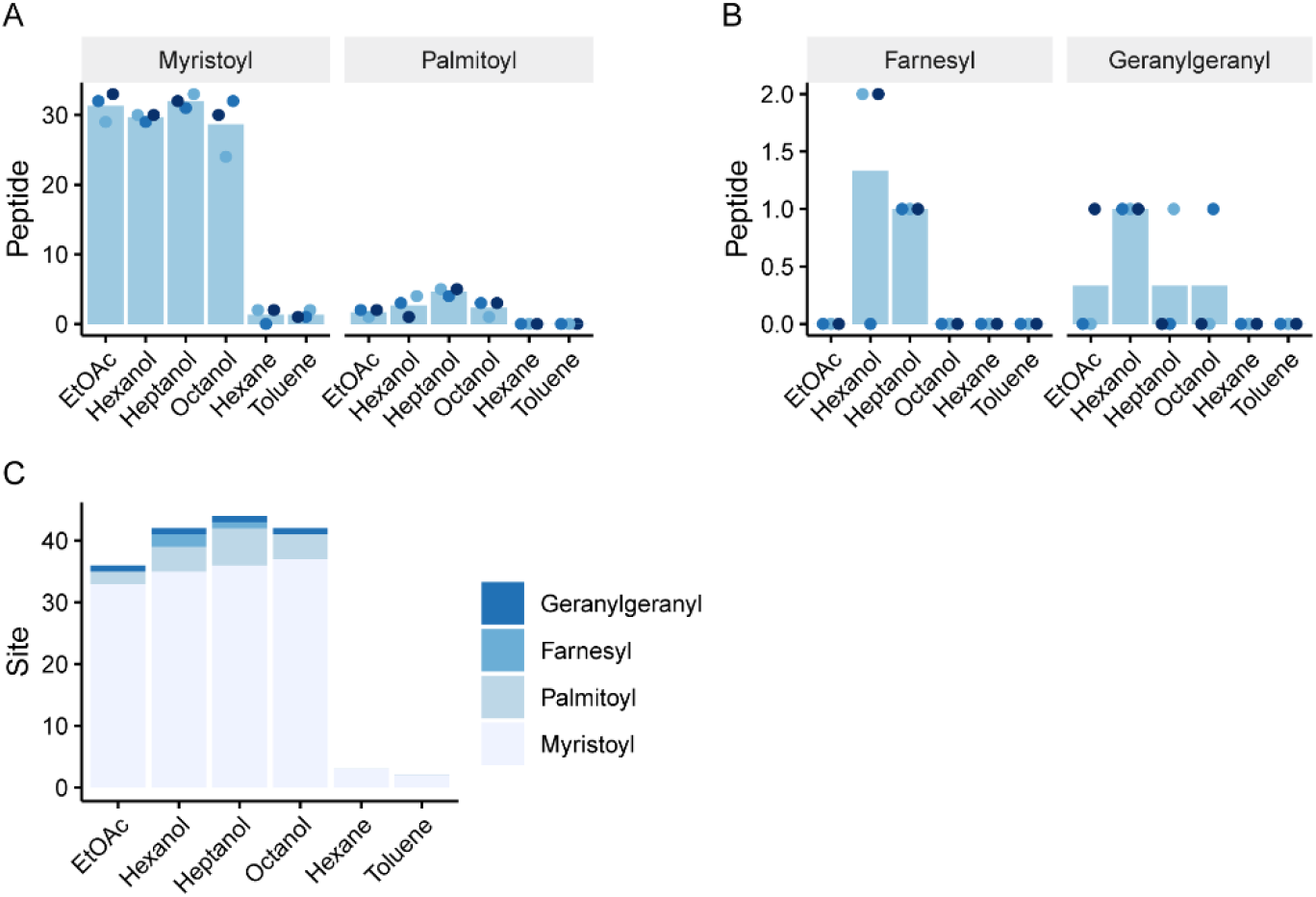
Comparison of organic solvents for extraction of lipidated peptides. (A, B) Numbers of identified myristoylated peptides and palmitoylated peptides (A) and farnesylated peptides and geranylgeranylated peptides (B). The results of triplicate experiments are indicated in different colors. Bars indicate mean values. (C) Summary of the numbers of identified lipidation sites. The results of triplicate experiments are combined. EtOAc, ethyl acetate.

### Lipidation profiling of HeLa cells

As a proof-of-concept of this methodology, we performed a comprehensive profiling of lipidation sites in HeLa cells. To expand the coverage, proteins were digested with trypsin/LysC, GluC, or chymotrypsin, in parallel (Fig. 3A). Aliquots of 100 μg of digests were subjected to LLE, in which heptanol and octanol were employed; thus, a total of 600 μg peptides was used per replicate. We first evaluated the results focusing on protein N-terminal myristoylation, for which the largest number of modified sites was identified among lipidations of interest in this study. The myristoylation sites were reproducibly identified in the technical quadruplicates (Fig. 3B). We identified a total of 75 myristoylation sites, which is greater than the number of the high-confidence lipidated proteins in one of the largest current protein N-terminal myristoylome datasets^10^ (Fig. 3C). The use of multiple digestive enzymes further increased the number of identified myristoylation sites (Fig. 3D). The sequences flanking modified glycine in our dataset are shown as a sequence logo in Fig. 3E, in which serine at the +5 position is highlighted. This motif is often observed in the case of protein N-terminal glycine myristoylation^22^, supporting the validity of our dataset. The identified sites included many known modification sites reported in UniProtKB, which contains 64 sites, including guanine nucleotide-binding protein G(i) subunit alpha (GNAI3; Fig. S4A) and ADP-ribosylation factor 1 (ARF1; Fig. S4B). We also found 11 sites not reported in UniProtKB, including plasminogen receptor KT (PLGRKT; Fig. S4C), indicating the ability of our methodology to identify unknown lipidated proteins at the site level. In addition, 9 palmitoylation sites were identified, of which four sites were reported in UniProtKB, including guanine nucleotide-binding protein G(s) subunit alpha isoforms short (GNAS) (Fig. S4D,). Interestingly, we detected three peptides dually modified with myristoylation and palmitoylation, derived from tyrosine-protein kinase YES (Fig. 3G), GNAI3 (Fig. S4E), and raftlin (RFTN1; Fig. S4F), all of which were reported in UniProtKB. Simultaneous observation of such multiple modifications is a major advantage of our methodology, since it is difficult to obtain information about modified sites by chemical proteomic approaches using lipid-mimetic probes. Moreover, four farnesylation sites, including peroxisomal biogenesis factor 19 (PEX19; Fig. S4G,), and two geranylgeranylation sites, including guanine nucleotide-binding protein G(i)/G(s)/G(o) subunit gamma-12 (GNG12; Fig. S4H), were identified. Four of these prenylation sites were reported in UniProtKB.

**Figure 3.**
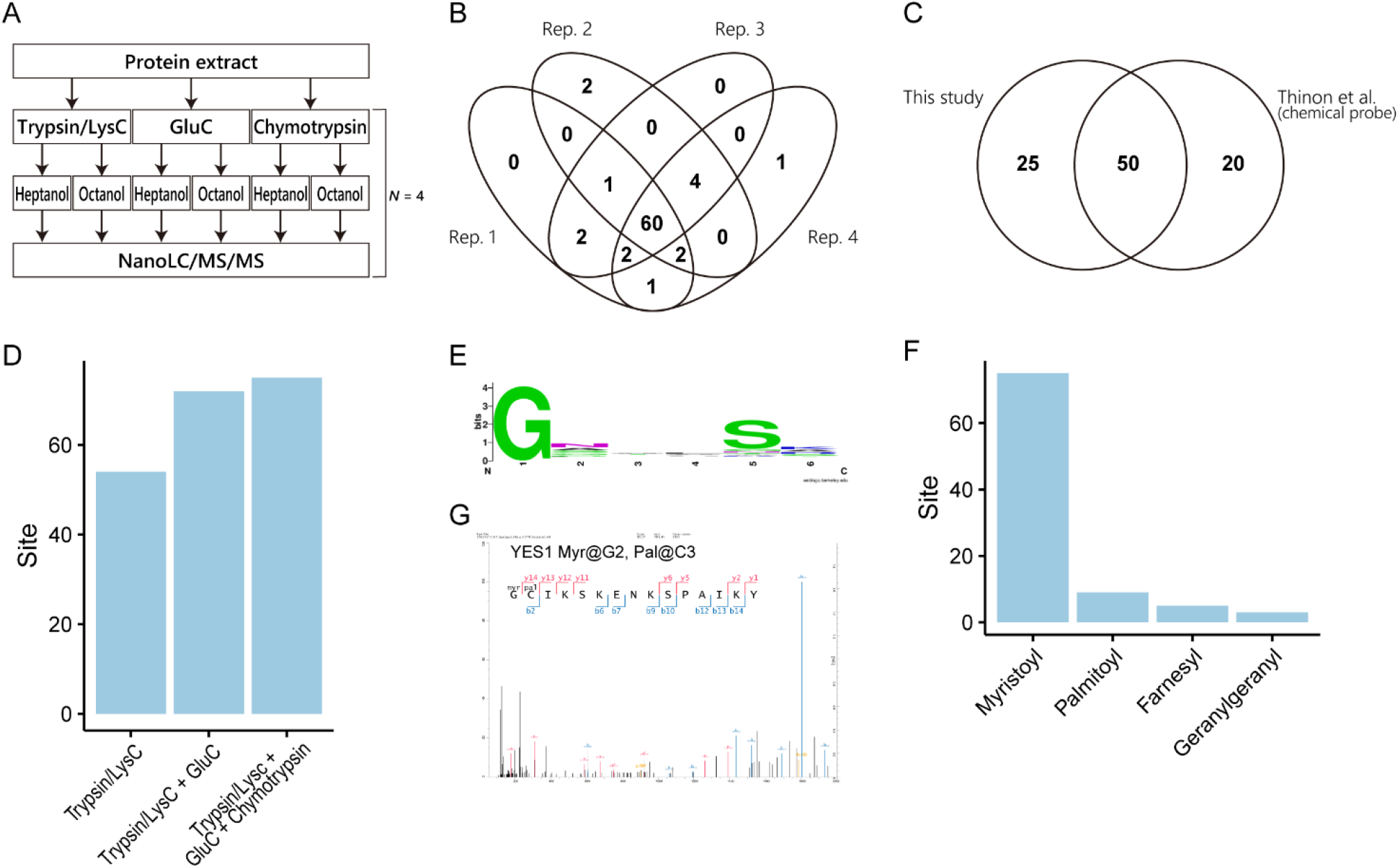
Lipidation profiling of HeLa cell proteins. (A) Overview of the workflow. From the same HeLa cell extract, three digests were prepared using different proteases. From each digest, lipidated peptides were isolated using heptanol or octanol in parallel and analyzed by nanoLC/MS/MS, which was done in quadruplicate. (B) Overlap of identified myristoylation sites among the technical quadruplicates. (C) Comparison of the identified myristoylation sites with the results of the largest current study on myristoylation. (D) Improvement in the number of identified myristoylation sites by combining the results obtained with different digestive enzymes. (E) Sequence logo showing amino acids downstream of the identified protein N-terminal myristoylation sites. (F) Summary of identified lipidation sites.

Compared to myristoylation sites, fewer sites were found for other lipidations, including palmitoylation. We assume that one of the reasons is the low stoichiometry of the modified forms. Co-translational protein N-terminal myristoylation would proceed at a high stoichiometric level during translation, whereas cysteine palmitoylation is dynamic^11^ and probably exists at a low stoichiometric level under basal conditions. Also, prenylation is known to accelerate degradation of some of its substrates^23,24^, which would make them more difficult to detect. Possible approaches to overcome these issues and facilitate deep analysis of low-stoichiometry modification sites might include the addition of a further processing step, such as subcellular fractionation and peptide fractionation by strong cation exchange chromatography (SCX) or the addition of inhibitors of the demodifying enzymes.

### Identification of *in vivo* lipidation sites in mouse tissues

Finally, we demonstrate that our methodology enables quantitative analysis of *in vivo* myristoylation sites in mouse tissues. We prepared protein tryptic digests from six mouse organs (kidney, muscle, colon, spleen, lung, and liver), and lipidated peptides were isolated using heptanol in three technical replicates. A total of 58 myristoylation sites were identified (Fig. S5). Among them, 23 sites, such as NIBAN1 and ABL2, were identified in all organs, while 8 sites, such as formin-like protein 3 (FMNL3) and HID1, were uniquely identified in a single organ (Fig. S5). The samples were segregated based on organs by hierarchical clustering (Fig. 4A). The Pearson correlation coefficients of log intensities of myristoylated peptides between the technical replicates in each organ were more than 0.98, whereas those between different tissues were lower (Fig. 4B). Taken together, our results indicate that this methodology is applicable for the identification of *in vivo* modification sites and is sufficiently reproducible to permit differential analysis.

**Figure 4.**
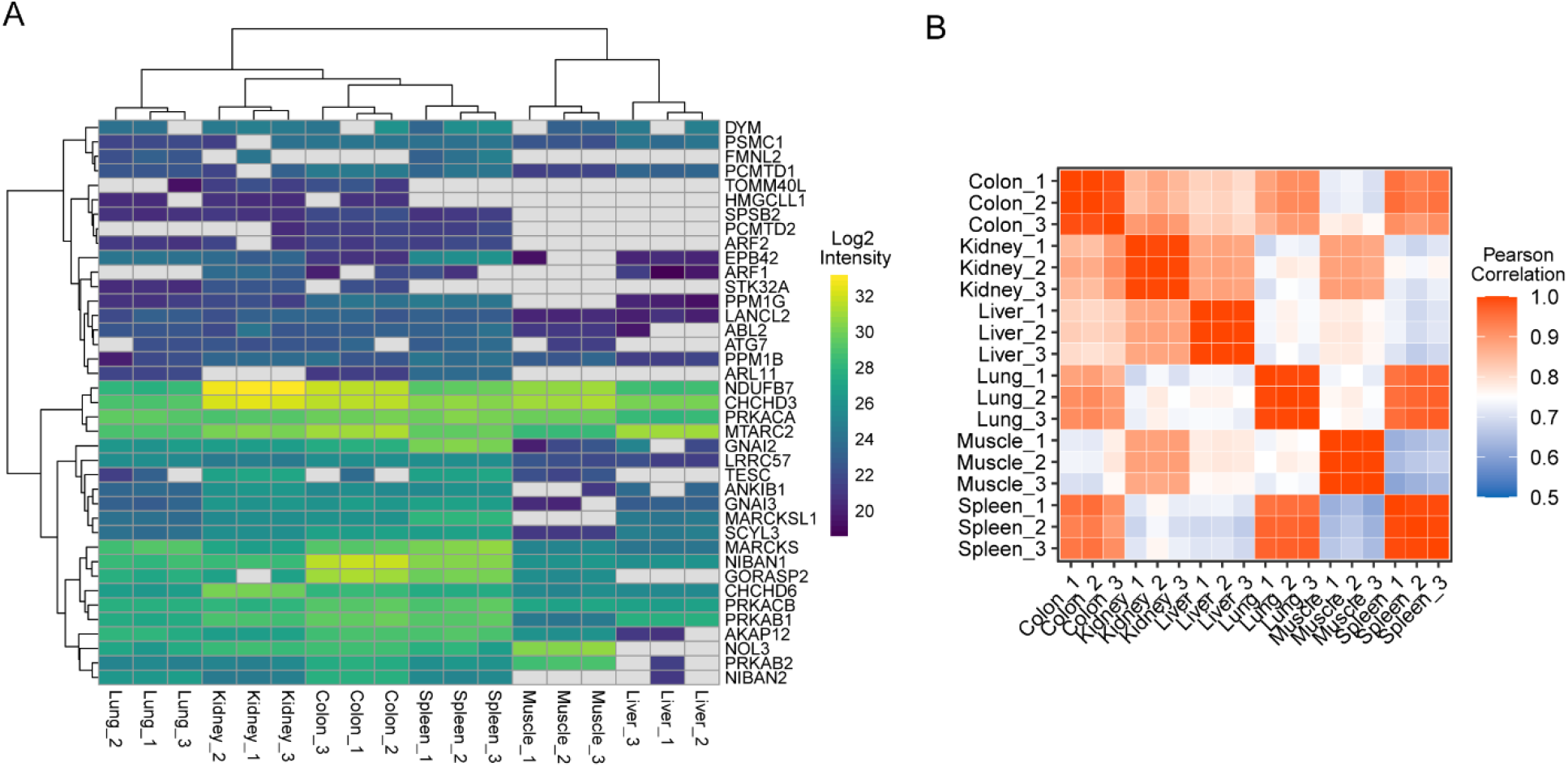
Comparison of myristoylation sites among mouse tissues. (A) Myristoylation sites in six mouse tissues are compared in technical triplicates. Note that the sites were filtered to missing value <20% for clustering, reducing the number of sites to 39. (B) Pearson correlation coefficients of the intensities.

## Conclusions

We have demonstrated for the first time that peptides modified with at least four forms of lipidation, myristoylation, palmitoylation, farnesylation, and geranylgeranylation, can be isolated by LLE, allowing direct identification by nanoLC/MS/MS using ACN in a higher concentration range. Our method does not require the cells to incorporate exogenous chemical probes, and is therefore applicable to intact biological samples. For low-stoichiometry modification sites, the addition of a fractionation procedure such as subcellular fractionation and peptide fractionation by SCX is expected to facilitate deeper proteomic analysis of lipidated peptides. This simple and rapid method should be useful to investigate biological machineries regulated by lipidation of functional proteins.

## Acknowledgments

This work was supported by Japan Science and Technology (JST) ERATO Arita Lipidome Atlas Project (grant no.: JPMJER2101 to KT, YIsobe, KI and MA), JST FOREST (grant no.: JPMJFR214L to KI), JSPS Grant-in-Aid for Scientific Research (Grant Number 20H03241 to KI) and RIKEN Special Postdoctoral Researcher Program (to KT).

## Supporting Information

Figure S1. Comparison of retention times between HeLa protein digest and synthetic lipidated peptides.

Figure S2. Representative MS/MS spectra of lipidated peptides, related to Figure 2.

Figure S3. Comparison of the number of myristoylation sites identified by different methods.

Figure S4. Representative MS/MS spectra of lipidated peptides in HeLa cells, related to Figure 3.

Figure S5. Protein N-terminal myristoylation sites detected in mouse organs.

## Notes

### Competing Interest Statement

The authors have declared no competing interest.

