## Supporting information for "Liquid-liquid extraction of lipidated peptides for direct identification of lipidation sites"

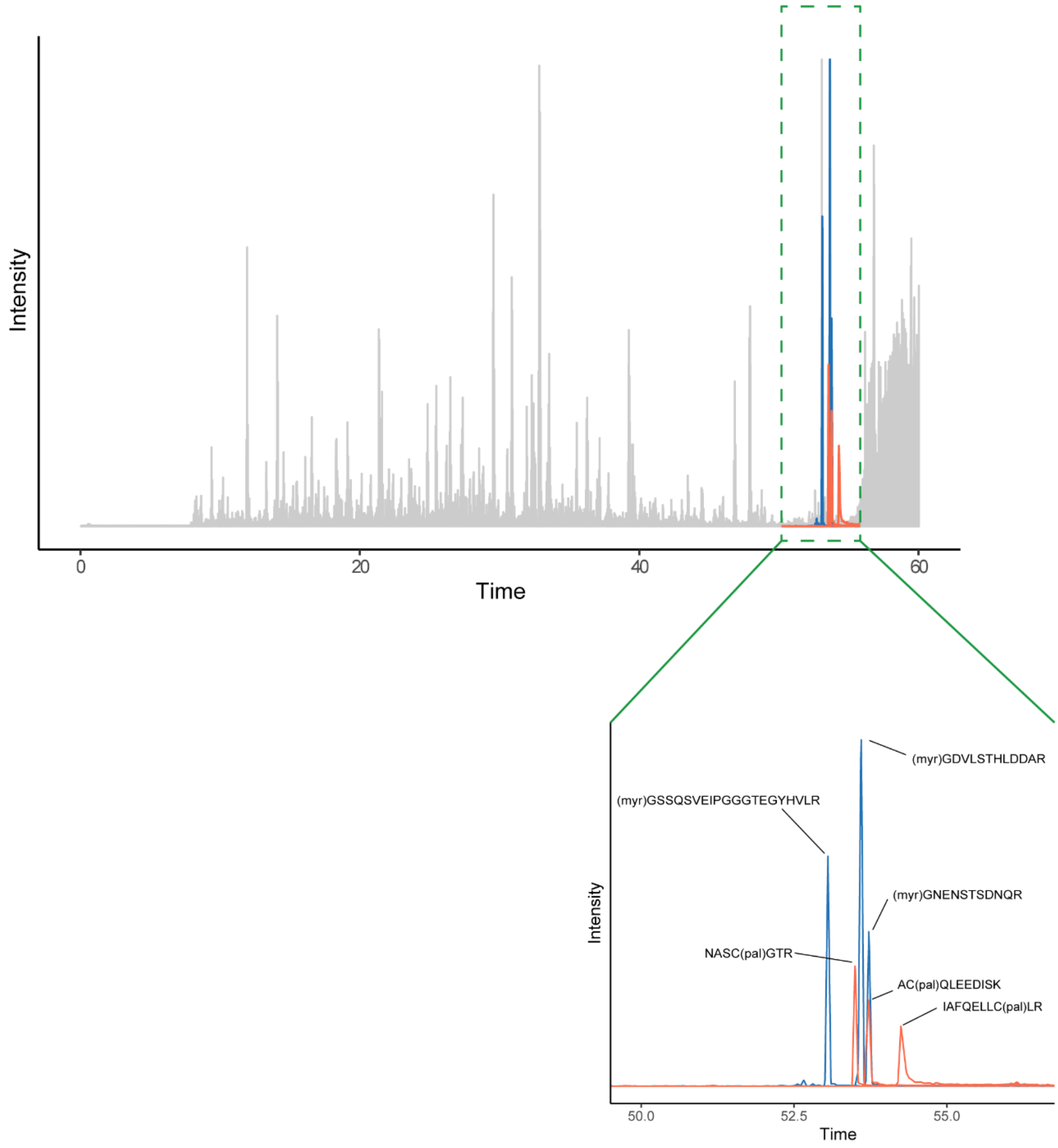


**Figure S1. Comparison of retention times of HeLa protein digest and synthetic lipidated peptides.**

Synthetic myristoylated peptides and palmitoylated peptides were spiked into HeLa protein digests and analyzed by nanoLC/MS/MS. The extracted ion chromatograms of myristoylated peptides (blue) and palmitoylated peptides (orange) are overlayed on the base-peak chromatogram. Note that intensities are normalized for ease of visualization.

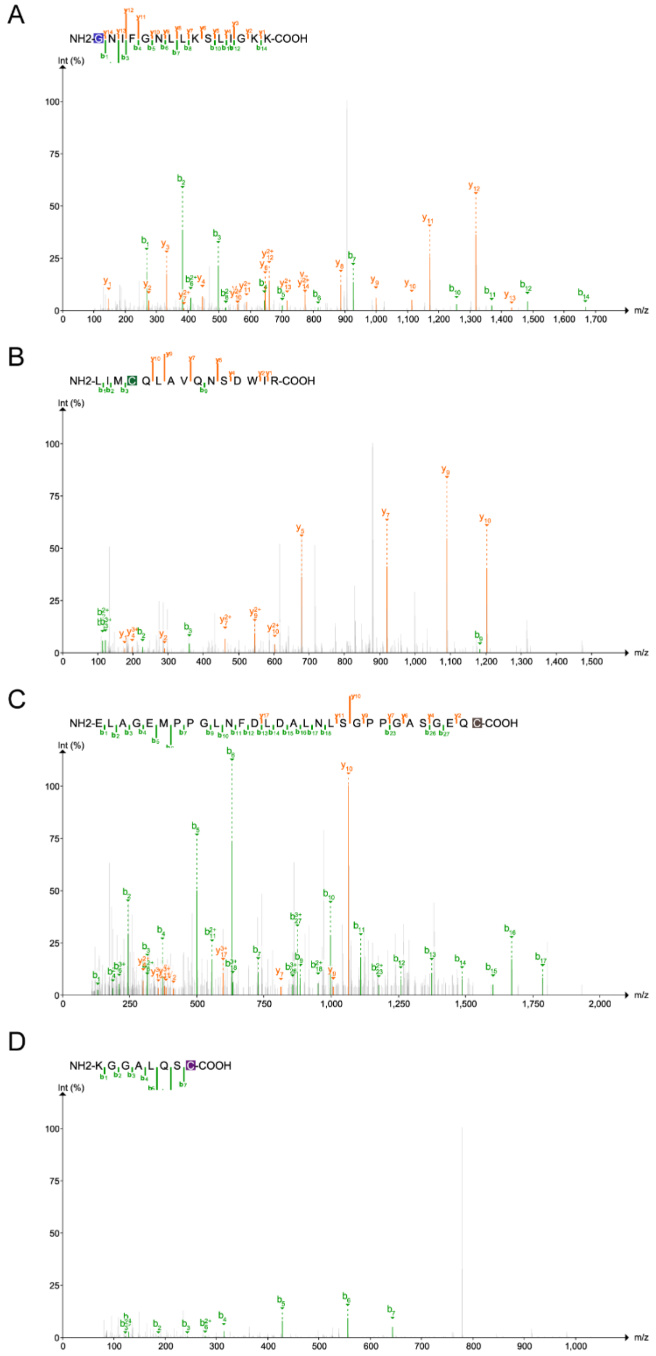


**Figure S2. Representative MS/MS spectra of lipidated peptides, related to Figure 2.** Representative MS/MS spectra of myristoylated peptides (A), palmitoylated peptide (B), farnesylated peptide (C), and geranylgeranylated peptide (D). (A) ARF3 G2; (B) NMNAT2 C70; (C) PEX19 C296; (D) CNP C418.

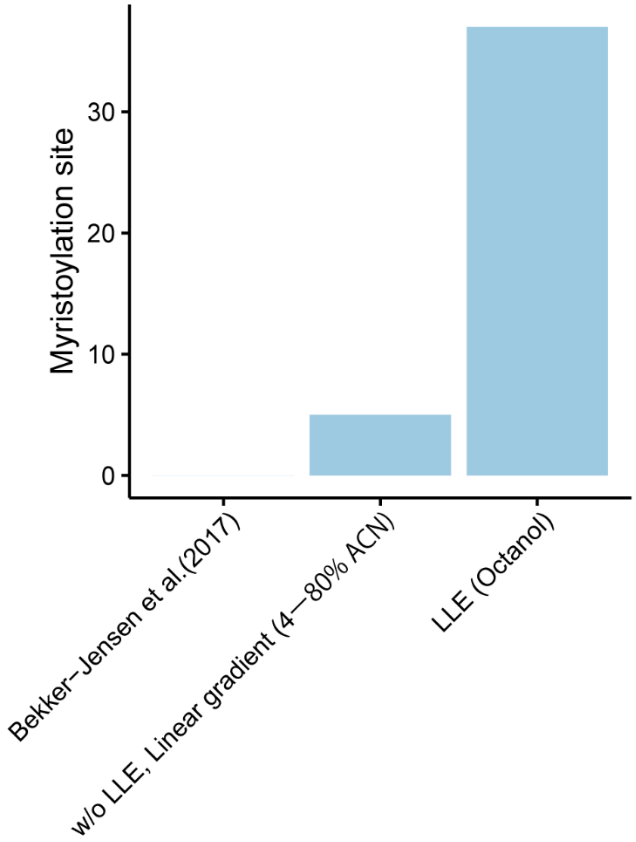


**Figure S3. Comparison of the numbers of myristoylation sites identified by different methods.** The number of identified myristoylation sites identified by liquid-liquid extraction using octanol (Fig. 2C) is compared with the result of re-analyzing the deepest current dataset of the HeLa cell proteome (Bekker-Jensen et al., *Cell Syst*. 2017) and the result of analyzing 500 ng of HeLa cell digest (identical to the input of LLE in this study) using a linear gradient ranging from 4 to 80 % ACN, without LLE. No myristoylation sites were found in the re-analysis of the dataset obtained by Bekker-Jensen et al.

*Supporting information*

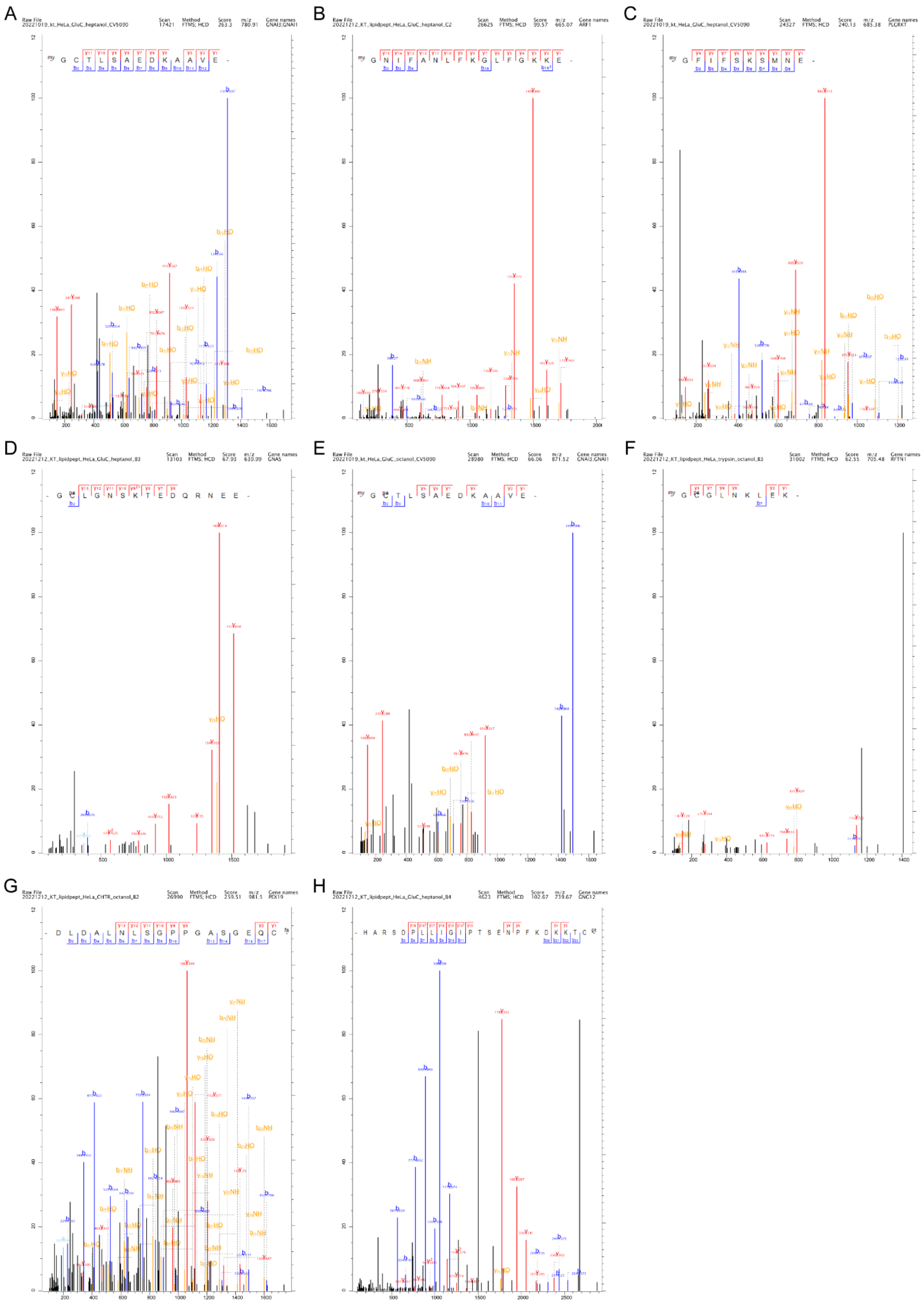

**Figure S4. Representative MS/MS spectra of lipidated peptides in HeLa cells, related to Figure 3.**

Representative MS/MS spectra of myristoylated peptides (A-C), palmitoylated peptide (D), peptide dually modified with myristoylation and palmitoylation (E, F), farnesylated peptide (G), and geranylgeranylated peptide (H). (A) GNAI3 G2; (B) ARF1 G2; (C) PLGRKT G2; (D) GNAS C3; (E) GNAI3 G2, C3; (F) RFTN1 G2, C3; (G) PEX19 C269; (H) GNG12 C69.

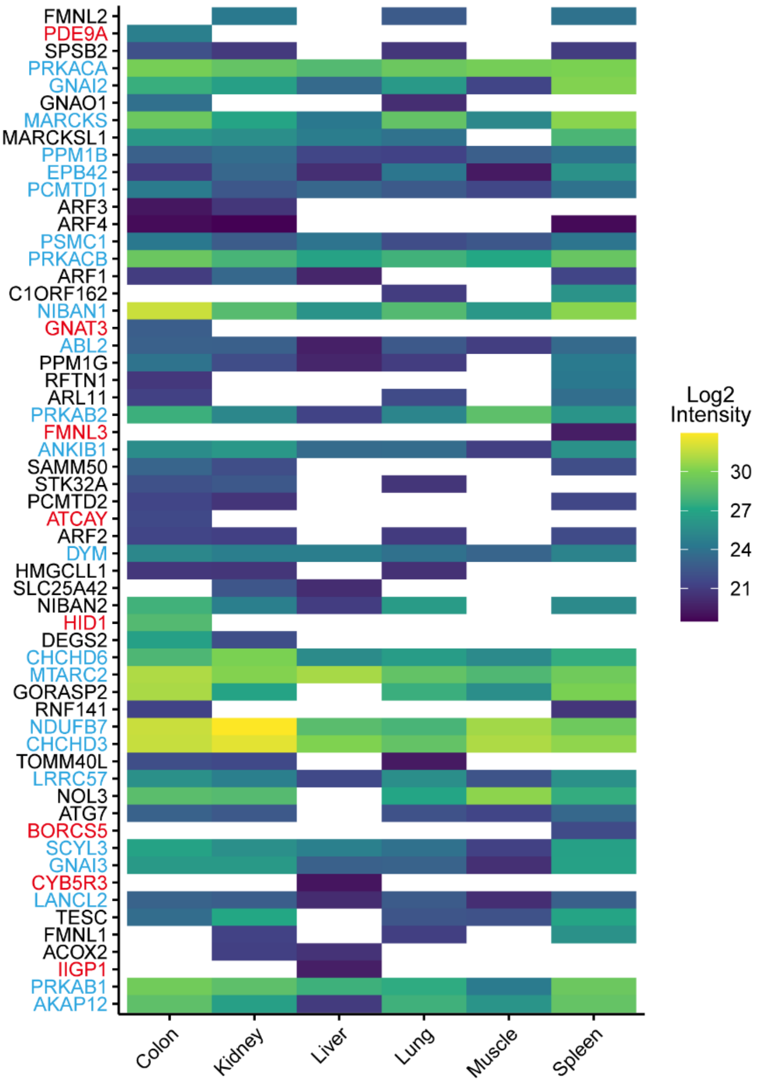


**Figure S5. Protein N-terminal myristoylation sites detected in mouse organs.**

Myristoylation sites identified in six mouse organs. Mean values of technical triplicates are shown. The sites uniquely identified in one organ and the sites commonly identified in all organs are highlighted in red and blue, respectively.
